# A Closed-Form Adaptation of the Hodgkin-Huxley Membrane Potential and its Synthesis to a Novel Electric Circuit

**DOI:** 10.1101/2022.07.25.501272

**Authors:** Robert F. Melendy, Loan Nguyen

**Affiliations:** Oregon Institute of Technology

**Keywords:** Action potential, axon, core conductor theory, cable theory, neuron, propagated signaling, field-dependent current, Hodgkin, Huxley, Langevin, intracellular magnetization, membrane conductance, membrane depolarization, membrane electric field, myelinated nerve, neuronal cable theory, electric network synthesis, membrane circuit, membrane capacitance, FitzHuhg-Nagumo nerve model, Na^+^ and K^+^ conductances, permeability efficacy

## Abstract

In a succession of journal papers published over 65 years ago, Sir Alan Lloyd Hodgkin and Sir Andrew Fielding Huxley discovered what now forms our contemporary understanding of excitation in nerve, and how axons conduct the action potential. Hodgkin and Huxley demonstrated that the nerve action potential is the result of a depolarizing event across a cell membrane. In an elegant theoretical framework, they established that when this depolarization event is complete, an abrupt increase in voltage gets produced that propagates longitudinally along the axon, accompanied by changes in axial conductance. Notwithstanding the elegance of Hodgkin and Huxley’s incisive and explicative series of discoveries, their model is relatively complex, relies on no small number of stochastic factors, and has no analytical solution; solving for the membrane action potential and the ionic currents requires integrations approximated using numerical methods. In this paper, we present a closed-form adaptation of the Hodgkin-Huxley membrane voltage potential. The basis of our model is rooted in core conductor theory and the cable properties of neurons, with fitting parameters adapted to the classical Hodgkin-Huxley model of excitation in nerve. From this model we synthesize a novel analog circuit that simulates the dynamics of a single action potential bioelectrically equivalent to the classical Hodgkin-Huxley membrane potential. The primary novelty of our model is that it offers a bioconductive, thermodynamic, and electromagnetic explanation of how an action potential propagates in nerve in a single mathematical construct. This is in contrast to the traditional Hodgkin-Huxley equations of ionic hypothesis, which are not analytically compliant. Computational results of our model are supported by well-established quantitative descriptions of Hodgkin-Huxley’s voltage response in the membrane of an axon. Our findings provide a mechanistic understanding of how intracellular conductance, the thermodynamics of magnetization, and current modulation function together to generate excitation in nerve in a unified closed-form description. In the same manner with Hodgkin-Huxley’s findings, the model presented here corroborates (1) that the action potential is the result of a depolarizing event across a cell membrane; (2) that a complete depolarization event is followed by an abrupt increase in voltage that propagates longitudinally along the axon; (3) that the latter is accompanied by a considerable increase in membrane conductance. The work presented in this paper provides compelling evidence that three basic factors contribute to the propagated signaling in the membrane of an axon in a single, closed-form model. From our model, we synthesize a novel analog conductance-level circuit that simulates the dynamics of a single action potential bioelectrically equivalent to the classical Hodgkin-Huxley membrane potential. It’s anticipated this work will compel those in biophysics, physical biology, and in the computational neurosciences to probe deeper into the classical and quantum features of membrane magnetization and signaling. Furthermore, it’s hoped that subsequent investigations of this sort will be advanced by the computational features of this model without having to resort to numerical methods of analysis.

**Attribution:** A portion of this work is reprinted from R.F. Melendy, <u>Resolving the biophysics of axon transmembrane polarization in a single closed-form description [1]</u>. *Journal of Applied Physics*, **118(24)**, Copyright © (2015); and from R.F. Melendy, <u>A subsequent closed-form description of propagated signaling phenomena in the membrane of an axon [2]</u>. *AIP Advances*, **6(5)**, Copyright © (2016), with the permission of <u>AIP Publishing</u>. Said published works are copyright protected by Robert. F. Melendy, Ph.D., and the AIP journals in which these articles appear. Under §107 of the Copyright Act of 1976, allowance is made for “fair use” for purposes such as criticism, comment, news reporting, teaching, scholarship, and research. Fair use is a use permitted by copyright statute that might otherwise be infringing. Nonprofit, educational (i.e., teaching, scholarship, and research) or personal use tips the balance in favor of fair use.

**Summary:** This work provides evidence that three basic factors contribute to propagated signaling in the membrane of an axon. The contributing factors are unified in a closed-form description. From this closed-form model we synthesize a novel analog circuit that simulates the dynamics of a single action potential that is bioelectrically equivalent to the classical Hodgkin-Huxley membrane potential.

## I. Overview and Scope

There is abundant well-grounded research quantifying the electrical behavior of myelinated and unmyelinated nerve fibers [3-6]. This has come to include an understanding of membrane impedance properties, and the longitudinal voltage and current signals that propagate in axon membranes [7-10]. Of note is Hodgkin and Huxley’s quantification of ionic membrane currents and their relation to conductance and excitation in nerve [11]. Since this time, more than a few researchers have focused on rigorously describing membrane phenomena and structure. A fundamental advancement in this direction was the development and prolific use of cable theory in modeling the membrane of an axon [12-16]. In classic cable theory, axons are treated as core conducting cylinders of finite length, where the capacitive and conductance properties of the axon membrane are modeled as a distributed-parameter electric network [17,18]. Consequently, determination of the membrane action potential and ionic currents requires the solution of a boundary-value problem. This approach provides a systematic means for realistically describing the action potential and the axon membrane field properties [19,20]. Nevertheless, this method of modeling repeatedly depends on the use of numerical methods to solve the partial differential equations. Comparably, the Hodgkin-Huxley equations of ionic hypothesis are a relatively complex system of differential equations that have no analytical solution; solving for the membrane action potential and the ionic currents requires integrations approximated using numerical methods.

The scope of this article is to derive an original, quantitative description of the membrane potential *V*_*m*_. From this description, we will synthesize an electric circuit that simulates the dynamics of a single action potential and that is bioelectrically equivalent to the classical Hodgkin-Huxley membrane potential. The order in which to accomplish this will be: (1) to present evidence that three principal factors form a basis on which the displacement of the membrane potential (i.e., from its resting value of ≈ –70 mV) is described; (2) to synthesize these factors into a single, closed-form expression for analytically computing *V*_*m*_; (3) to demonstrate the range of phenomena to which the mathematical form is relevant. The latter will be achieved by: (*a*) substitution of established membrane parameters into the mathematical form, followed by; (*b*) computation of the membrane electric field, *E*_*m*_; (*c*) computation of *V*_*m*_ from its resting value through the hyperpolarizing afterpotential. Computational results will be compared with classical standards. It is from this model that we will synthesize an electric circuit whose response is electrically equivalent to a single action potential.

## II. Synthesis of the Membrane Potential Analytical Model

In this section, both electrodynamic and thermodynamic evidence will be presented in forming a basis on which the displacement of the membrane potential *V*_*m*_ is described. These phenomena – in due course – will be presented in a unified, analytical description of membrane excitability, followed by a description of *V*_*m*_ in terms of a single, nonlinear, homogeneous differential equation. The basis of the analytical model will be established in neuronal cable theory. Only certain features resulting from cable theory are of relevance to the development of the analytical model. Accordingly, the applicable research will be sufficiently referenced.

### A. The Leaky Cable Conductance Property of an Axon

One solution to the neuronal cable equations is a function describing the input resistance *R*_*in*_ (Ω) of a leaky cable along the longitudinal length of neuronal fiber [21,22]:

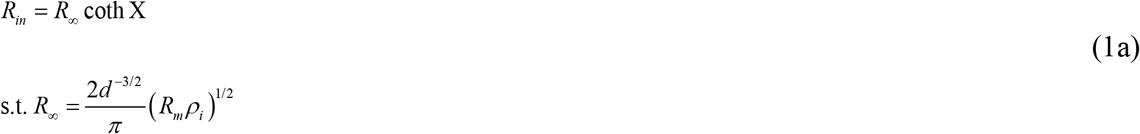

where *R*_∞_ is the input resistance of a semi-infinite cable and is proportional to the characteristic length, *λ* (m) of a membrane cylinder. *R*_*m*_ represents the resistance across a unit area of membrane (Ω·cm^2^), *ρ*_*©*_ is the resistivity of the intracellular medium (Ω·cm), and *d* is the diameter of the membrane cylinder (∼μm). The property of intracellular resistivity is related to the axoplasmatic resistance to movement of electric charge *q* © [23,24]. Extracellular resistance is considered negligible. X (Chi) is a normalized length (dimensionless). Normalized length is often given the notation “*L*” in the literature, but this is too easily confused with an actual (physical) length (m). In this article, X is used in place of *L*. A non-ideal X will not be constant but will vary along the physical length *x* of the axon [18,23]. It is defined by X = ∫ 1/*λ dx* for cylindrical membranes. This is integrated over the distances along successive (compartmental) cylindrical axes.

It is elementary to rewrite the hyperbolic cotangent of (1a) as *R*_*in*_ = *R*_∞_ (1/tanh X) or *R*_∞_ (cosh X/sinh X). This is identical to writing (*R*_*in*_/cosh X) = (*R*_∞_/sinh X) ⟹ (*R*_*in*_/cosh X) = (*R*_∞_ csch X). Since resistance is the reciprocal of conductance *G* (Ω^−1^), the left-hand term may be expressed as (1/*G*_*in*_ cosh X). From this simple arrangement of terms, one can write:

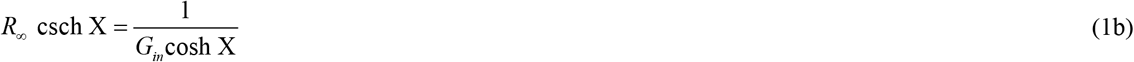

The biophysical relevance of (1b) is it describes how a rapid drop in the input resistance of a semi-infinite cable (*R*_∞_) balances with a significant decrease in the leaky cable input resistance (*R*_*in*_) along the longitudinal length of neuronal fiber.

It’s traditional and convenient to express the movement of ionic charge and polarization changes in the membrane of an axon in terms of conductance. By and of itself, the hyperbolic conductance term (1b) is intrinsic to the displacement of the membrane potential, *V*_*m*_ (V) from its resting value:

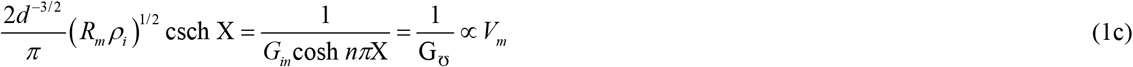

The inverse variation (1c) is consistent with the fact that voltage varies inversely with conductance [25]. For initial generality, *nπ* multiples of X are initially included in the cosh argument. The left-hand units of (1c) is ohms (Ω). From Ohm’s law, (1c) is consistent with the fact that *V* ∝ *R*.

### B. Axon Intracellular Magnetization Hypothesis

When a membrane depolarizes, a natural consequence is the generation of a changing magnetic field. There’s a body of established research corroborating the existence of time-varying magnetic fields in an axon during the nerve impulse [26-30].

A common thread that runs through these studies is that the bioelectric activity present during the action potential produces a current in a volume conductor. For instance, the current density *J* (A·m^−2^) throughout a volume conductor generates a biomagnetic field, *B* (T). Without exception, the latter exists in axon membranes and have been experimentally shown to be of remarkably small magnitude [27,28]. These studies offer a classical description of biomagnetic field phenomena.

In contrast, what can be understood about the biomagnetic field of an axon membrane from a statistical mechanics description? Could such a description be unified with the macroscopic conductance term (1c)?

Biological tissue has been shown to have paramagnetic properties, particularly in the presence of Ca^+^ and Na^+^ ions [31-33]. It is therefore relevant to consider intracellular magnetization as an intrinsic membrane property (particularly, during the action potential event). Langevin’s paramagnetic equation is suitable in this circumstance: (*M* /*μN*) = tanh (*μB*/*kT*) [34]. Where *M* is magnetization (A·m^−1^ or J·T^−1^·m^−3^), *N* is the number of particles that make up the membrane material [with each particle having magnetic moment *μ* (J·T^−1^)], *k* is Boltzmann’s constant (1.38 × 10^−23^ J·K^−1^), and *T* is temperature (K).

Langevin’s equation predicts that a paramagnetic material saturates asymptotically to the line (*M* /*μN*) as (*μB*/*kT*) → 2 [35]. In this instance, the fast-microscopic variables are the statistical averages of the noise generated by the thermal fluctuation of electrons in the conducting axon. The thermodynamic derivation of this noise predicts the electrical response of the axon to the resting and membrane response potentials quantified by the conductance. During polarization for instance, there’s a considerable increase in the sodium conductance *g*_Na_ of the membrane. This produces a marked increase in the current density throughout the conducting medium [36] and subsequently, an appreciable increase in the magnetization of the intracellular membrane. By Langevin’s relation, it stands to reason that this intracellular magnetization saturates as all the moments become aligned against the biomagnetic field during a complete polarization event.

On the basis of this hypothesis, the hyperbolic conductance term (1c) and Langevin’s thermodynamic relation are asserted to vary together, according to:

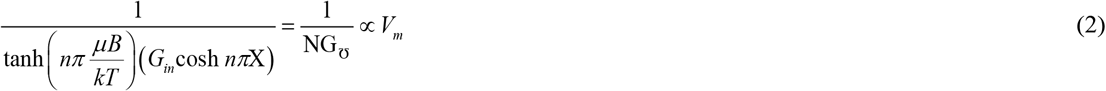

Observe that (2) has left-hand units of Ω (i.e., since the hyperbolic tangent term is dimensionless). As with (1c), *nπ* multiples of (*μB*/*kT*) are initially included in the hyperbolic argument for generality. From Ohm’s law, (2) is consistent with the fact that *V* ∝ *R*.

### C. The Membrane Current Modulation Hypothesis

An accepted and reliable method for depolarizing the excitable cells of a membrane involves variations in voltage-clamping techniques [37,38]. Regardless of method, the sensors utilized in voltage-clamping exploit the properties of the membrane potential and ionic current signals [39]. These signals are not fundamental. They’re constructed of sinusoidally-varying harmonics of the form *A* cos *ω t, B* sin *ω t*, or some convolution of these functions. Some signals have been shown to be unstable depending on the initial conditions in the membrane [40,41]. Irrespective of harmonics or stability, the usual practice is to quantify these signals as functions of time. The same holds true for the description of biomagnetic signals in an axon. This raised the question: Can the membrane current that accompanies the action potential be quantified in terms of the biomagnetic field, i.e., *I* = *I*(*B*)?

#### 1. Field-Dependent Current Premise

One can deduce *a priori* that a current *I*(*B*) inevitably propagates through an axon of physical length *l* for the period of a depolarizing event. This is perfectly reasonable since time-varying magnetic fields have been measured in axon membranes during depolarization and hyperpolarization (as previously discusses and referenced on p.4). By Ampere’s law, a field-dependent current *I*(*B*) must therefore exist, such that ∮ *B* · *dl* = *μ*_0_ *I*(*B*).

This prompts the question: Can one quantify variations in *I*(B) for the period of a depolarizing event? This would suggest the presence of a *current modulation signal, d* ^2^ *I*(*B*)/*dB*^2^ (A·T ^−2^). As is characteristic of the classically understood Na^+^ and K^+^ time-dependent currents, it’s reasonable to assert *I*(*B*) would also exhibit non-fundamental oscillatory-like behavior. It is well understood that a membrane response to a depolarization current pulse is accompanied by a rapid drop in the leaky cable resistance (and hence, a net increase in intracellular conductance). This necessitates a marked rate of increase in *I*(*B*) during depolarization (as discovered and referenced by Roth and Wikswo, p.4) [27].

To mathematically synthesize a function for *I*(*B*) (and to demonstrate the range to which its mathematical form is relevant), the cylindrical geometry of a classic axon [23-25] and its intrinsic electromagnetic behavior are considered basic. This is a perfectly reasonable deduction, and lends itself to physical problems involving cylindrical coordinates. For instance, the description of electromagnetic fields in cavities (e.g., field strength behavior far-from and close-to cavity walls) [42,43] makes use of spherical Bessel functions, *j*_*n*_(*x*). In series notation, the spherical Bessel function is written *j*_*n*_(*x*) = (–1)^*n*^ *x*^*n*^ (*x*^−1^ *d* /*dx*)^*n*^ (sin *x*)/ *x*, where *n* is an integer (0, 1, 2, 3,…, *n*).

##### Axiom

The existence of a field-dependent current *I*(*B*) induced in the membrane of an axon must be a response to some input excitation. Even if this excitation were an ideal impulse *δ* (*x*), the membrane could never produce a *δ* response (this would be physically impossible, and no experimental results have ever shown this to be the case). This would necessitate that the series sin (*x* / *l*) /*π x* → *δ* (*x*) in the limit as the axon length *l* → 0 (physically impracticable). Since *l* can never → 0 in the limit, it follows that *I*(*B*) must consists of a finite number of quantitative terms. If *I*(*B*) is therefore to be modeled by a collection of spherical Bessel functions, then by the arguments made here, *I*(*B*) would consist of only the first few integer values of *n* [44].

Neurons of membranes have been shown to have natural frequency-selective feedback properties [45,46]. It stands to reason that such properties would influence how the field-strength current *I*(*B*) gets transmitted, absorbed, reflected, etc., based on frequency during the action potential. This seems particularly true when one considers the observation of close to subcritical Hopf bifurcations in neurons, with membrane conductances and currents functioning as bifurcation parameters [41,47,48]. These phenomena support the presence of the current modulation signal, *d* ^2^ *I*(*B*)/*dB*^2^.

### 2. Field-Dependent Signal Convolution Postulate

On the premise of the preceding discussion, it’s reasonable to expect that *d* ^2^ *I* (*B*)/*dB*^2^ would exhibit fluctuations through the membrane for the period of a depolarizing event. This is supported by the elementary fact that a magnetic field cannot instantaneously collapse in an axon as the action potential transitions from depolarization to the hyperpolarizing afterpotential. One plausible conjecture is that the current modulation signal would behave according to *d* ^2^ *I*(*B*)/*dB*^2^ ∝ *f* (*B*) ⊗ *j*_*n*_(*B*), where *f* (*B*) is some induced electromagnetic response signal and ⊗ is convolution. For now, it must be postulated that *f* (*B*) is not constant but varies nonlinearly in response to *B*(*t*). Furthermore, the magnetic field must be of relatively adequate strength such that the membrane energy density (J·m^−3^) is sufficient to completely depolarize the membrane.

### 3. Synthesis of the Current Modulation Function Postulate

There are chaotic nonlinearities associated with initiation of the nerve impulse by membrane depolarization [49,50]. This, taken in conjunction with the oscillatory nature of the spherical Bessel functions *j*_*n*_(*x*) (particularly, for *n* = 0 to 2), one could reasonably hypothesize that *d* ^2^ *I*(*B*)/*dB*^2^ will exhibit unstable oscillations for the period of a depolarizing event [51-53].

Without exception, unstable eigenvalues are nearly always present in dynamic systems. In biological systems, there are intrinsic control mechanisms that operate in the presence of unstable equilibrium points to produce a stable response, often *after* a margin of instability [40,54-56]. On the premise of unstable oscillations, the simplest of cases would be a signal quantified by the Bessel function (–1)^*n*^ *x*^*n*^ (*x*^−1^ *d*/*dx*)^*n*^ sin *x*/*x* multiplied with *f* (*B*) = *x*^2^ for *n* = 0, yielding *x*^2^ *j*_0_ (*x*) = *x* sin *x*. The initial prediction is therefore a modulation signal of the form *d* ^2^ *I*(*B*)/*dB*^2^ = (2*π a* /*μ*_0_) × (*t* sin *nπω t*), where the inclusion of (2*π a* /*μ*_0_) is a consequence of Ampere’s law, *a* being the axon radius (∼μm), and *μ*_0_ the vacuum permeability (4*π* × 10^−7^ H·m^−1^).

### 4. An Initial Quantitative Description of the Action Potential

The question posed in this section (p.4) – *Can the membrane current that accompanies the action potential be quantified in terms of the biomagnetic field, i*.*e*., *such that I = I(B)?* – can now be addressed. By Ohm’s law, *V* ∝ *I*. Hence, the displacement of the membrane potential *V*_*m*_ must be in proportional variation to *I*(*B*) and any *n*^th^ derivative of *I*(*B*), such that:

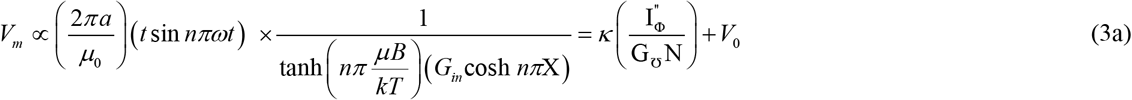

where I_Φ_^″^ = (2*π a*/*μ*_0_) × (*t* sin *nπω t*). The intracellular resting potential of the membrane *V*_0_ (i.e., relative to the outside of the cell) must also be accounted for. As before, *nπ* multiples are initially included in the transcendental for computational generality.

The term 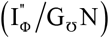 has units of V·T^−2^. In order to reduce this term to units of volts, a constant of proportionality *κ* having units of T^2^ is introduced. A postulate is that *κ* = the square of the membrane magnetic field *B*_*m*_ (T), such that:

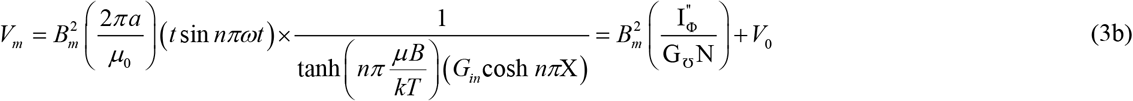

The numerator 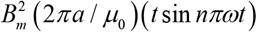 of (3b) is a current term having units of amps (A). The denominator of (3b) is conductance (Ω^−1^). From Ohm’s law, (3b) is consistent with the fact that *V* = *IR*. (3b) offers an initial analytical description of the membrane action potential *V*_*m*_. As per the scope of this article (p.2), the next step will be to rewrite (3b) in terms of the accompanying cell membrane electric field, *E*_*m*_.

## III. The Membrane Electric Field Hypothesis

Consider once again the current term 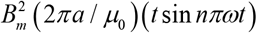 in (3b). Electric current *I* in the axon per unit area of the axon cross section is *J* = *I*/*A* (A·m^−2^), where *A* = *π a*^2^ and *a* is the axon radius. The current density may also be written as *J* = *E*_*m*_ /*ρ*_*m*_, where *E*_*m*_ is the axon membrane electric field (V·m^−1^) and *ρ*_*m*_ is the longitudinal membrane resistivity (Ω·cm or Ω·m). Hence, the axon current flow may be expressed as *I* = *JA* = (*π a*^2^)*E*_*m*_ /*ρ*_*m*_. By the laws of electrodynamics [57], the numerator of (3b) may therefore be re-written as 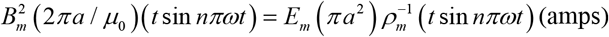.

The electric field is constant along the axon longitudinal axis but is radial dependent, such that **E**_*m*_ = *E*_*m*_ **û**_*r*_, where **û**_*r*_ is a unit vector in the axon radial direction. It’s more practical therefore to express 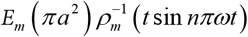 in terms of the axon thickness, Δ*r* [18,23-25] (i.e., such that *E*_*m*_ **û**_*r*_ ∞ Δ*rE*_*m*_). Consider the introduction of the proportionality constant *k*, such that *k*Δ*rE*_*m*_ produces units of amps (A). Then *k* would need to have units of (F·m^−1^) × (V·m^−1^)^2^. The conjecture therefore is that *k* = *ε*_0_ *E*_*m*_, where *ε*_0_ is the vacuum permittivity of free space (8.854 × 10^−12^ F·m^−1^). Substituting these relations into (3b) gives:

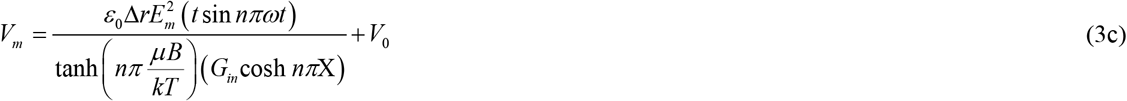

The units of volts are preserved in going from (3b) to (3c). It will be subsequently shown that (3c) gives a correct description of the classic nerve action potential and the cell membrane electric field. Computational results for *V*_*m*_ and *E*_*m*_ will be validated by comparison with standardized values in the literature.

## IV. Materials and Methods

A `Matlab` algorithm was developed to computationally test the biophysical adaptation of the model (3c). This required a practical choice of membrane physical parameters [14,23,25,35,36] (Table 1):

**Table 1:** The physical parameters used to compute the voltage potential (3c).

| Physical Parameters | Symbol and Value |
| --- | --- |
| *Axon thickness (myelinated-constant)* | $\Delta r = 2 \mu\text{m}$ |
| *Axon ("cable") length* | $0 \leq x \leq 4000 \mu\text{m}$ |
| *Length constant* | $\lambda = 1000 \mu\text{m}$ |
| *Resistance (unit area of membrane)* | $R_m = 2.56 \Omega \cdot \text{m}^2$ |
| *Intracellular resistivity* | $\rho_i = 0.4 \Omega \cdot \text{m}$ |
| *Input resistance (semi-infinite cable)* | $R_\infty = 20.3718(32) \text{ M}\Omega$ |
| *Nonlinear magnetization (unitless)* | $0 \leq (\mu B/kT) \leq 4$ |
| *Action potential cycle time* | $0 \leq t \leq 5 \text{ msec}$ |
| *Vacuum permittivity* | $\varepsilon_0 = 8.854(10^{-12}) \text{ F/m}$ |

But testing the accuracy of (3c) also requires gauging it against a scientifically accepted standard, namely, the numerically integrated membrane potential *V*_*m*_ from the Hodgkin-Huxley equations of ionic hypothesis [9].

Hodgkin and Huxley hypothesized that the net current *I*_*m*_ which flows into a unit area of membrane surface is the sum of the current *l*_*C*_ flowing into the membrane capacitance *C*_*m*_ (per unit area) and the ionic current *I*_*i*_ associated primarily with sodium and potassium species:

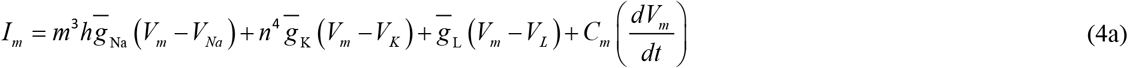

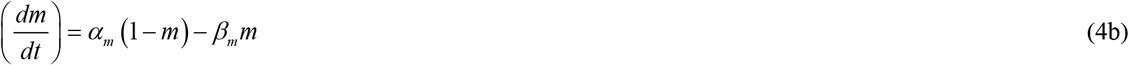

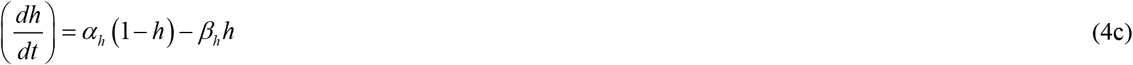

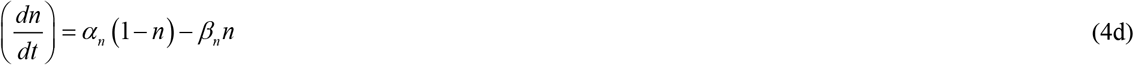

where *V*_*m*_ is the membrane action potential (mV). The potential of the sodium, potassium, and leakage channels are denoted by *V*_Na_, *V*_K_, and *V*_L_, respectively. The maximum conductance associated with each species are denoted by 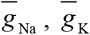, and 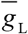, respectively (mmhos/ cm^2^). The controlling parameters *m, h*, and *n* are time-varying coefficients ∈ (0, 1) and represent the probability that any channel is open to the flow of ionic currents *I*_Na_ and *I*_K_ (m and hare associated with two types of sodium channels, whereas n is associated solely with potassium). Each *α* and *β* is an experimentally observed rate-constant derived from kinetic theory [5-7,9], and each is approximated by a smooth function of the membrane voltage *V*_*m*_. For the squid axon at a temperature of 6.3°C:

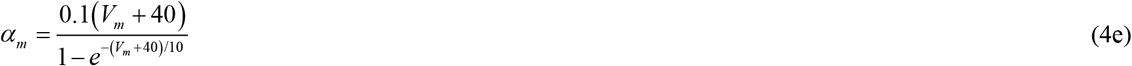

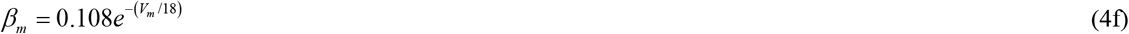

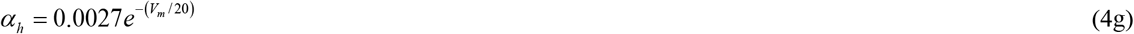

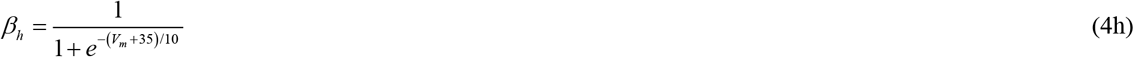

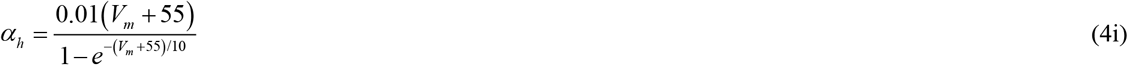

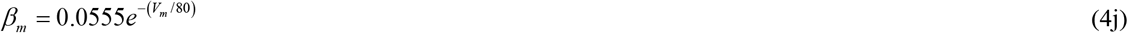

A `Matlab` code was written using a fourth-order Runge-Kutta algorithm to numerically integrate differential equations (4a) through (4d). Furthermore, the `Matlab` algorithm was coded to solve for the stochastic rate constants (4e) through (4j) [this was necessary to simultaneously integrate (4a) through (4d)]. The primary goal was to obtain a numerically integrated vector containing the data points for the action potential curve, *V*_*m*_.

In keeping consistent with Hodgkin and Huxley’s original experiment, we used standard published values of the membrane parameters [5,9] in our simulation (Table 2). Upon execution of our `Matlab` code, we successfully obtained a numerically integrated vector containing the data points for the action potential curve, *V*_*m*_. This enabled us to numerically fit (as close as possible) the latter to the unknown parameters of the adaptation model (3c). Namely, the integer values *n* in each of the transcendental arguments and the radial frequency term *ω* (radians/sec or s^−1^) in the numerator term of (3c).

**Table 2:**
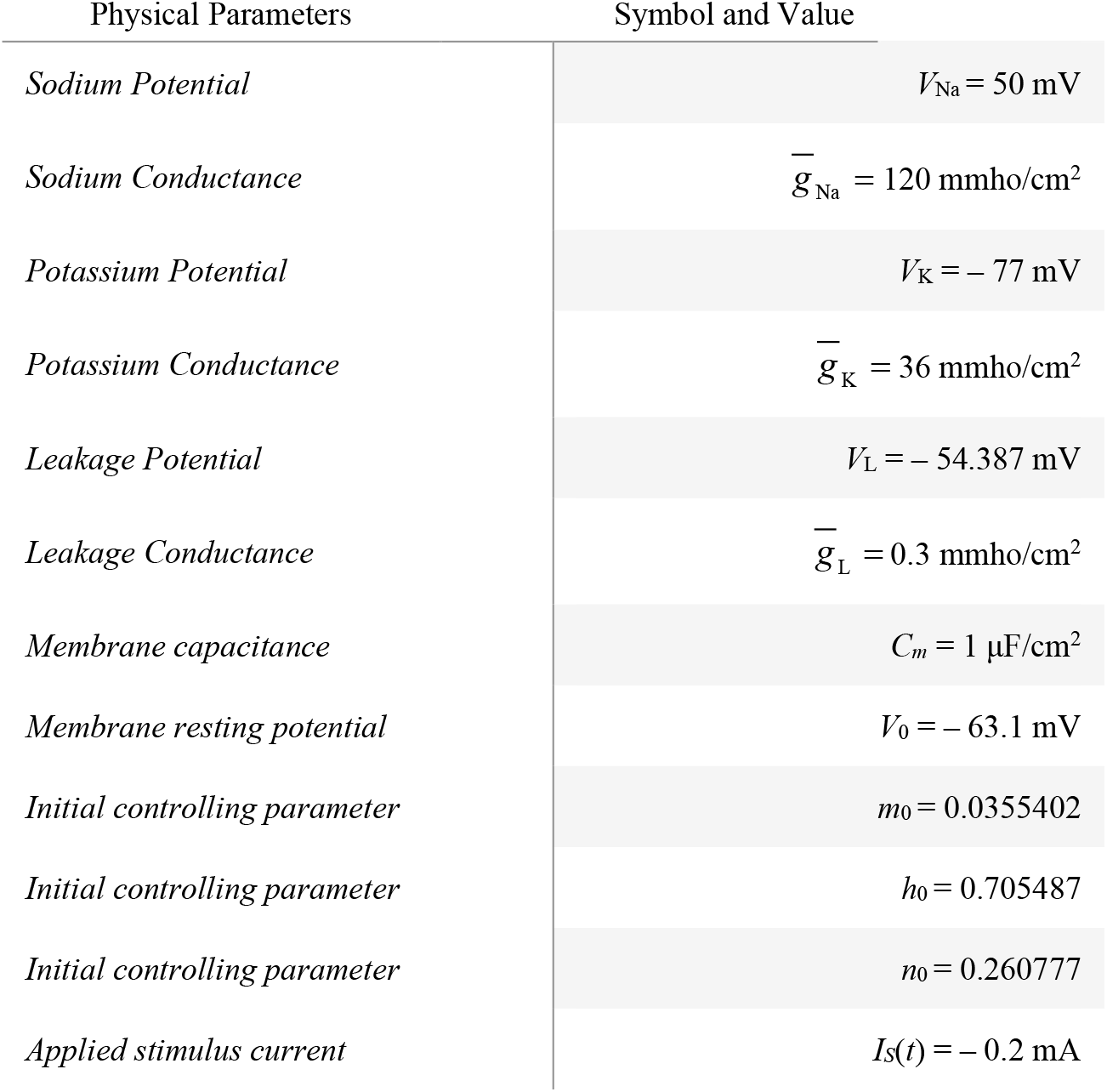
The physical parameters necessary to compute the classical Hodgkin-Huxley action potential (4a) through (4j).

For the cosh (conductance) argument in (3c), a best-fit iteration returned a value of *n* ≈ 1. The dimensionless magnetization factor tanh (*nπμB*/*kT*) of (3c) was not providing the parameterization necessary to correctly model the unique dynamics of a classical membrane action potential. An asymptotic series expansion [58] of this expression was performed to reveal the sensitivities associated with each of the terms in the tanh argument. It was found that this term was best-fit to an exponential function, such that *b*^tanh^(^*nπμB*/*kT*^).

In fitting our `Matlab` algorithm to the numerically integrated vector containing the data points for the Hodgkin-Huxley action potential (4a) through (4j), we discovered that *b* ≈ 0.475. For *n* in the tanh argument of (3c), the algorithm returned a best-fit iteration of *n* ≈ 4. In parameterizing the field current term 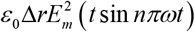, the `Matlab` algorithm returned a best-fit iteration of *n* ≈ 1 and *ω* ≈ ½ s^−1^ for the sin argument. However, the simulation kept resuming a somewhat abnormally-shaped action potential response. It was concluded that the hypothesized field term was not suitably parameterized based on postulates (*1*) through (*3*) (p. 4-6). This raised the question: Was this term displaying a sensitivity-dependence on the initial conditions? This reasoning supported the notion of *Lyapunov’s stability criterion*, and the possible need for computing a Lyapunov characteristic number, *ξ* [59]. The Lyapunov characteristic number provides information about the rate of separation of infinitesimally close trajectories. Classically, *λ* is used for the Lyapunov characteristic number, but *λ* has been used in this article for the axon length constant, so *ξ* was chosen. A double-precision floating point `Matlab` algorithm was written to compute *ξ* from the physical parameters chosen. This resulted in the estimate *ξ* = 2.704(77) ≈ *e*. The fact that *ξ* > 0 was not surprising since the field current signal was predicted to exhibit unstable oscillations during membrane polarization (p.6). This term was corrected to account for the sensitivity-dependence of the field current signal.

On the basis of these computations, the closed-form model (3c) can thus be amended:

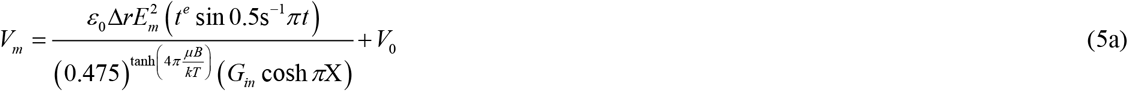

### A. Reliability of the Hypothesized Model (5a): Computation of the Membrane Electric Field, *E*_*m*_

Our `Matlab` algorithm was used to establish an estimate of the term 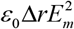 to initiate complete depolarization of the axon, converging to a value of 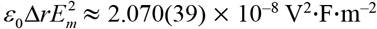. The average thickness of a myelinated cell membrane is Δ*r* ≈ 2 μm (Table 1) [23,60,61]. It follows that 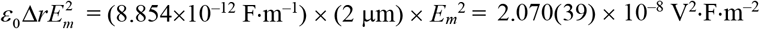. This results in *E*_*m*_ = 3.418(95) × 10^4^ V·m^−1^. A classic axon membrane model will have a potential difference between the interior and exterior side of the membrane of Δ*V*_*m*_ ≈ –70 mV [62]. The theoretical electric field for a myelinated membrane of 2 μm thickness is therefore *E*_*m*_ = – *dV*_*m*_/*d*(Δ*r*) = – (–70 mV)/(2 μm) = 3.5 × 10^4^ V·m^−1^ [23,60,63]. This result is favorably consistent with the computation from the hypothesized model (5a), having a percent error on the order of ≈ 2.3%. This is an initial confirmation that (5a) is a correct description of the classical membrane action potential, *V*_*m*_ [11]. In compact notation, we let 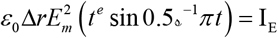.

### B. Reliability of the Hypothesized Model (5a): Computation of the Membrane Potential, *V*_*m*_

To further substantiate that (5a) is an equivalent adaptation of the nerve action potential established by Hodgkin and Huxley [11], a computational profile of *V*_*m*_ was completed for 0 ≤ *t* ≤ 5 msec. Compiling all preceding factors into the Matlab algorithm gives a restoration voltage *V*_0_ ≈ –70.0 mV. In completed form, (5a) is now expressed as:

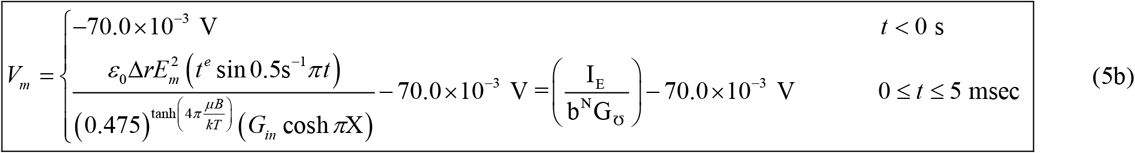

Fig. 1 is a plot of *V*_*m*_ vs. *t* from: (*i*) (4a)-(4j); (*ii*) (5b). Both plots demonstrates the classical action potential voltage signal in nerve under stable equilibrium conditions [11,18,23,25,36,61,63].

**Fig. 1.**
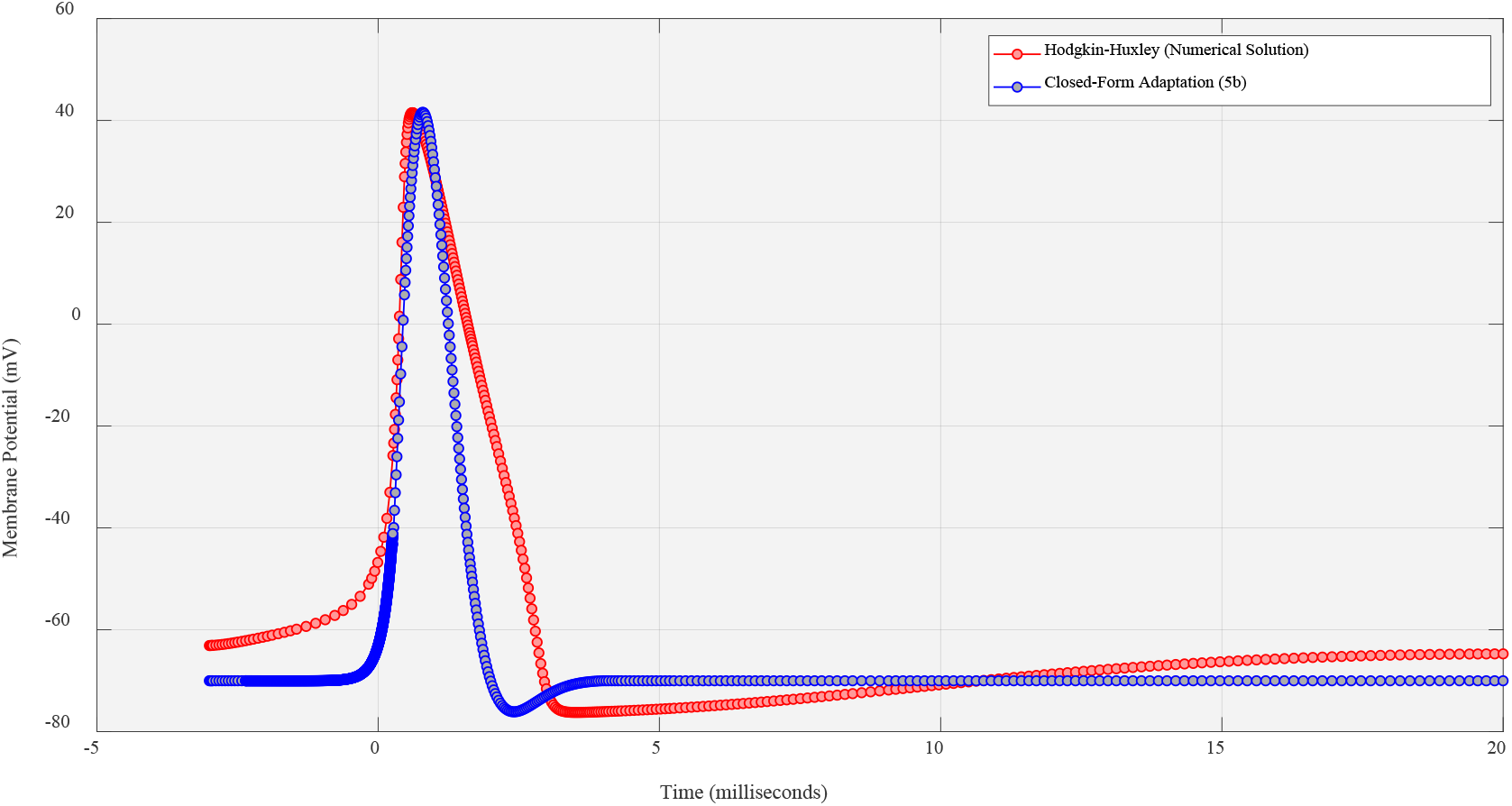
Comparison of Hodgkin-Huxley’s action potential from (4a)-(4j) and the closed-form adaptation of (5b).

## V. Discussion: Inference of the Ionic Current Flow

For the Hodgkin-Huxley equations of ionic hypothesis (4a)-(4j), it’s a well-established fact that the lipid bilayer of the axon membrane is modeled as a lumped-capacitance *C*_*m*_ (F) [11,64]. This concept is illustrated in Fig. 2. Also well-established is the quantification of the ionic current flow *I*_*C*_ through this bilayer, such that *I*_*C*_ = *C*_*m*_ × (*dV*_*m*_ /*dt*) [(as described in (4a)].

**Fig. 2.**
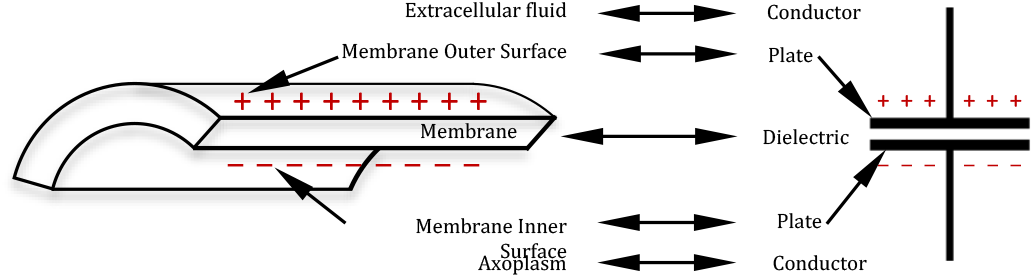
Neuronal membranes behave in part as if it were composed of a lumped capacitance. The membrane represents the dielectric while the extracellular fluid and the axoplasm represent the conductors.

The relationship between the membrane electric and magnetic fields *E*_*m*_ and *B*_*m*_, respectively, may be expressed in terms of the time rate-of-change of the membrane potential, such that *dV*_*m*_ /*dt* = *E*_*m*_^2^/ *B*_*m*_. The current flow through the membrane lipid bilayer may likewise be expressed in terms of these fields, such that: 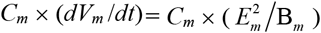. Thus, a novel feature of (5b) is that of offering an alternative description to the classical model for the time-dependence of the membrane current *I*_*C*_ in terms of *E*_*m*_.

The field current term 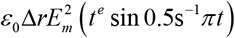 of (5b) has units of amps (A), where 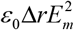 has units of V^2^·F·m^−2^ or T^2^. The point is that 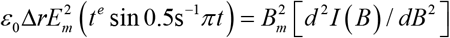, meaning that the time-dependent membrane current *I*_*C*_ of the Hodgkin-Huxley model (4a) is inferred by the electric field-induced current term implicit in (5b). It is hoped this makes clear the implicit manifestation of the membrane current underlying the action potential and its relationship to the membrane electric field, *E*_*m*_. The concluding hypothesis is that (5b) resolves the biophysics of how an axon conducts the action potential in a unified closed-form adaptation.

## VI. Design Methodology: Synthesis of (5b) to a Novel Electric Circuit

From Ohm’s law, (5c) is consistent with the fact that *V* = *IR*. For the time being, we ignore the restoration voltage *V*_0_ term. The circuit design will consist of three basic systems:

1. A circuit for the current modulation term: 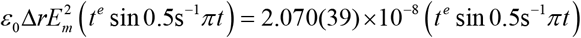
2. A circuit for the membrane resistance term: 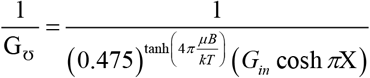
3. Multiplier, amplifier, and clamping circuits.

### A. The Conceptual Circuit from Matlab Data

The `Matlab` algorithm already exists to generate the necessary plots of the membrane current and resistance (Fig. 3). These plots are the basis for selecting and determining the topology and components of the circuits.

**Fig 3.**
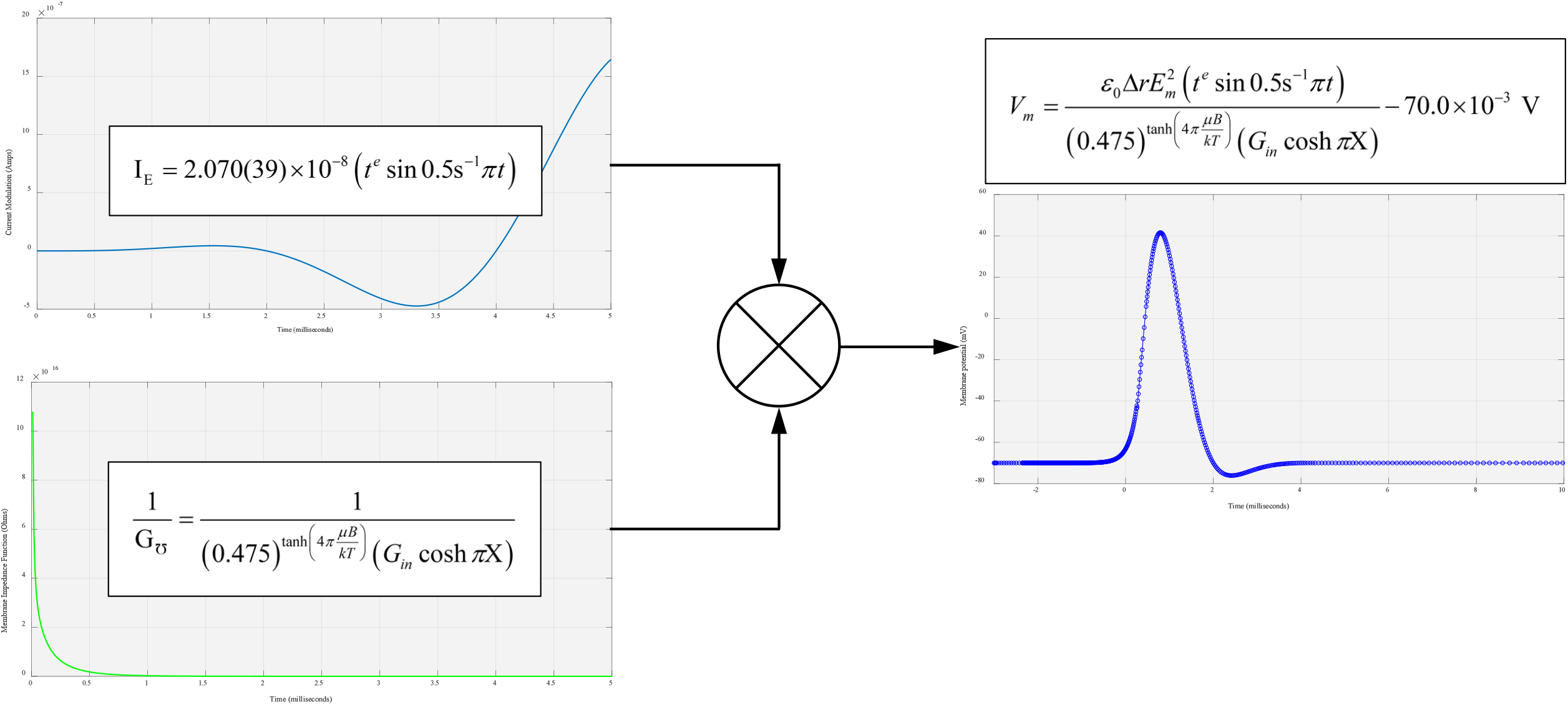
Conceptual diagram of circuit to produce the desired action potential response (5b) through signal convolution.

### B. Circuit Synthesis from the Current Modulation Function

For the time being, we will ignore the numerical value 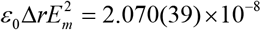 and focus our design efforts on the product *t*^*e*^ sin 0.5s^−1^*π t*. We will (1) synthesize two circuits corresponding to each function [65,66]; (2) use an AD633 integrated circuit [67] to perform a convolution operation on the output responses of the latter (Fig. 3). For the term sin 0.5s^−1^*π t* we apply a sine source to the input (pin #3). The remaining problem is for the signal *t*^*e*^. The value of *t*^*e*^ at *t* = 5 ms is 79.4347 ≈ 80. From this, we reasoned that the term 80*e*^−*t*^ can be rationalized as a function of a discharging capacitor voltage response with an initial (step) value of 80 V. Graphs of 80*e*^−*t*^ and *t*^*e*^ are nearly symmetrical about the vertical axis at *t* = 2.5 ms. We therefore considered the process of charging a capacitor to create the symmetric response *t*^*e*^. The growth rate of *t*^*e*^ however is marginally slower than the rate of decrease of its symmetric function 80*e*^−*t*^. This necessitated the addition of an inductor to our design and an adjustment of the excitation pulse such that the slope of the response fitted the slope of *t*^*e*^ during 5ms of the process (Table 3).

**Table 3.** Slopes of *t*^*e*^ and our charging capacitor prototype at various intervals of the action potential cycle.

| Interval to calculate slope | Slope of $t^e$ | Slope of the plot of charging capacitor |
| --- | --- | --- |
| (0.5-1) msec | 1.7 | 1.7 |
| (1-2) msec | 5.58 | 5.88 |
| (2-3) msec | 13.38 | 13.56 |
| (3-4) msec | 23.49 | 22.18 |

Our prototype is therefore predicted to be as a series *RLC* circuit having an initial capacitor voltage *v*_*C*_(0), an input excitation *u*(*t*), and an output response *v*_*C*_, such that:

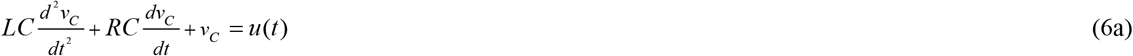

From the information in Table 3, we conclude that the time constant for the charging capacitor is *τ* _*C*_ ≈ 2.3s = *RC*. Considering for now the homogeneous response of our prototype, we let *u*(*t*) = 0 and take the Laplace transform of (6a). This gives the *s*-domain equation for the capacitor voltage:

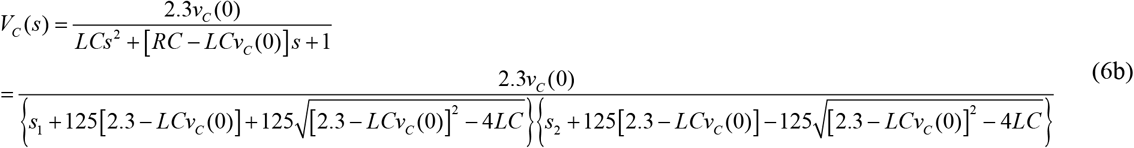

The complete response of our prototype circuit will be of the form:

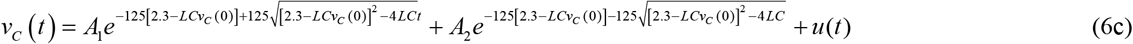

We ran a parameter sweep simulation in LTspice XVII [68] to identify a best-fit response slope of (6c) matching that of *t*^*e*^ during 5ms of the process. From this, our design values are *R* = 8Ω, *L* = 0.125H, and *C* = 0.29F (Fig. 4).

**Fig 4.**
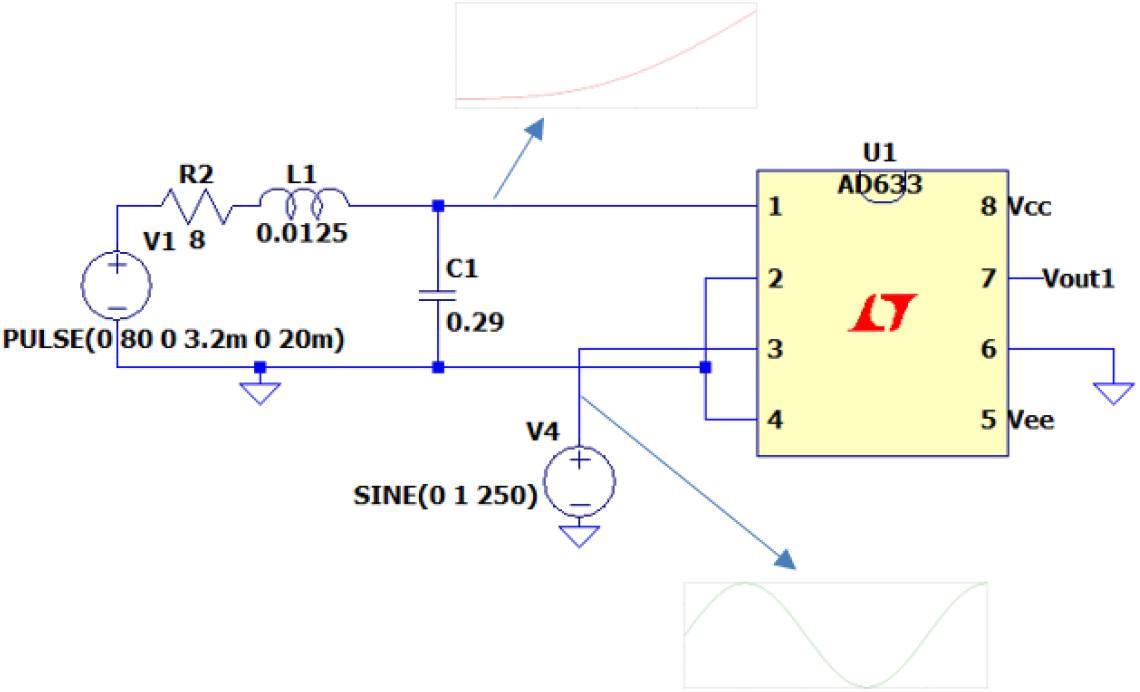
LTspice XVII simulation of *t*^*e*^. The signal Vout1 (pin #7) represents the `Matlab` modulation response of Fig. 3.

### C. Circuit Synthesis from Membrane Resistance Term

In Section B (p.13), we provisionally omitted the term 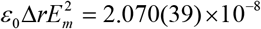. In order to synthesize a circuit having the same parametric scale as our preceding design (Fig. 4), we divide the membrane resistance term by 2.070(39) × 10^−8^. The resulting plot is illustrated in Fig. 3 (p.12) and at first glance, appears to be exponential. This was suspicious due to the extremely large values of 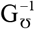 for very small *t*. We examined this distortion more closely by generating a semilog plot of 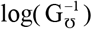 vs. time (Fig. 5). The slope of 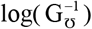 for the first 5ms is tabulated in Table 4.

**Table 4.** The slope of 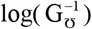 at various intervals of the action potential cycle.

| Time (milliseconds) | Resistance ($10^9 \Omega$) | Time Interval to Calculate Slope | Slope of $\log(R)$ |
| --- | --- | --- | --- |
| 1 | 234,700 | - | - |
| 2 | 8,027 | (1-2) msec | -2,266 |
| 3 | 336.1 | (2-3) msec | -7,690 |
| 4 | 14.46 | (3-4) msec | -321.64 |
| 5 | 0.6245 | (4-5) msec | -13.835 |

**Fig 5.**
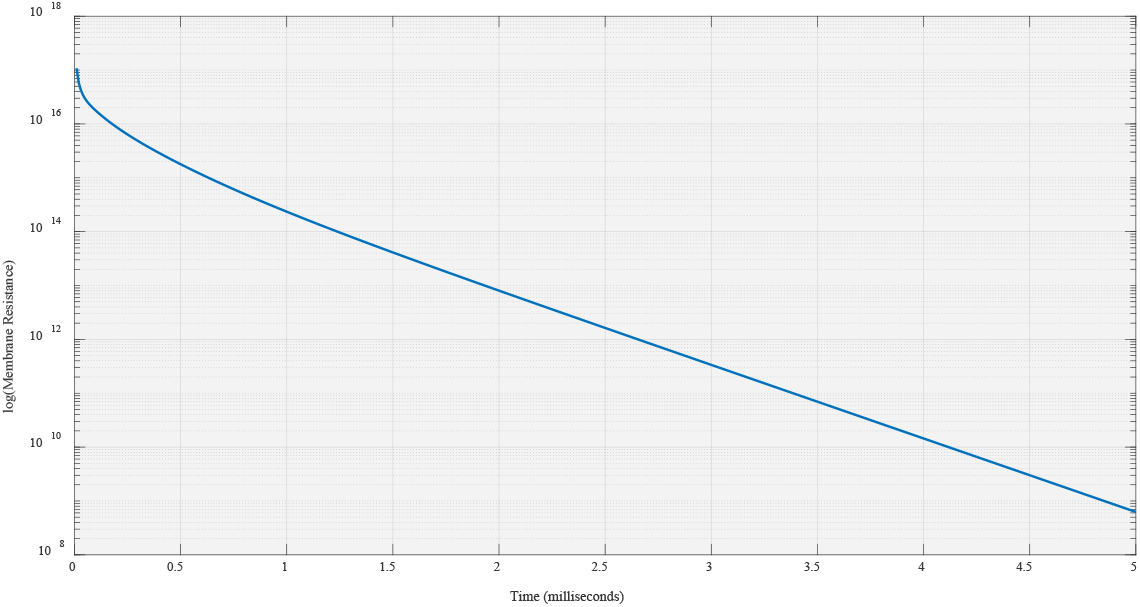
The plot of log(*R*) vs. time.

The process of discharging a capacitor was used to generate an output voltage corresponding to the behavior of membrane resistance as per Table 4. For the slope of a circuit response to coincide with the slope of the data in Table 4, we ran a parameter sweep simulation in LTspice XVII to identify a best-fit response slope coinciding with the slope of 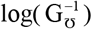 during 5 ms of the process. From this, we discovered that the time constant for a capacitor discharging through a 20 kΩ resistor is *τ* _*C*_ ≈ 0.0002 s, or *C* ≈ 0.01 μF. The decay rate of 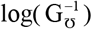 was slower than the rate of decrease of *RC* simulated circuit. So as previously done we added an inductor to shape the slope of the circuit response to match the slope of 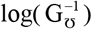 during 5 ms of the process. For our 20 kΩ resistor, we discovered that the time constant for the *RL* portion of the circuit is *τ* _*L*_ ≈ 0.001 s, or *L* ≈ 2 H. The diode *D*_1_ was included to force the capacitor *C* to discharge through the *RL* network (Fig. 6).

**Fig 6.**
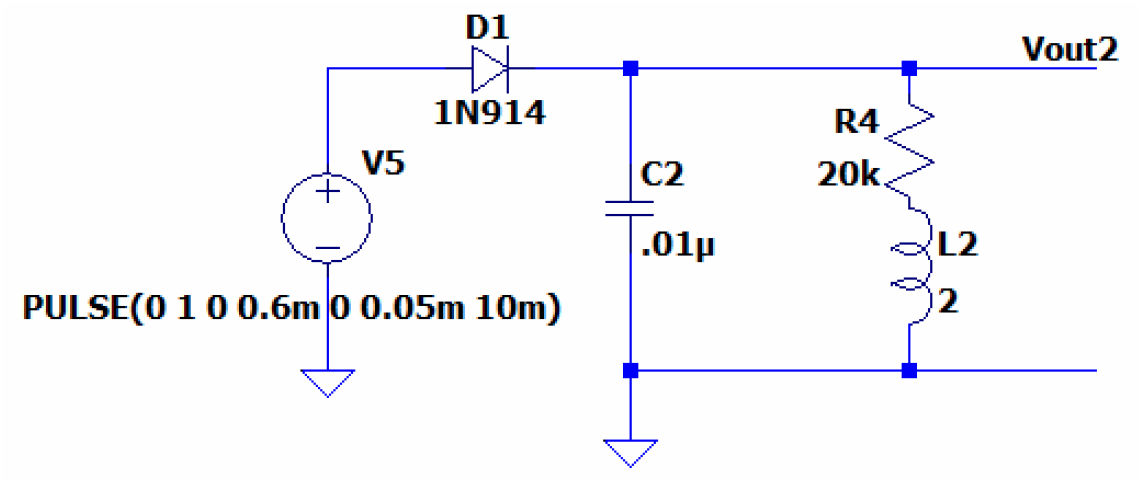
LTspice XVII simulation. The signal at Vout2 represents the `Matlab` resistance function of Fig. 3.

### D. Convolution of the Responses

The response of each of the filters (Fig. 4 and Fig. 5) must undergo a convolution process so as to produce a response that assimilates the action potential of (5b) (see Fig. 3). We again used an AD633 integrated circuit to perform this convolution operation on the signals Vout1 and Vout2 (Fig. 7). The analog multiplier connections are as follows: (1) input signal 1 comes from pin #7 of Fig. 4; (2) input signal 2 comes from the voltage response of Fig. 5; (3) the response of the second AD633 (pin #7) is the result of the convolution product (Vout1⊗Vout2)+voltage value of pin #6 (the voltage at pin #6 is used to shift the level of the output signal).

**Fig. 7.**
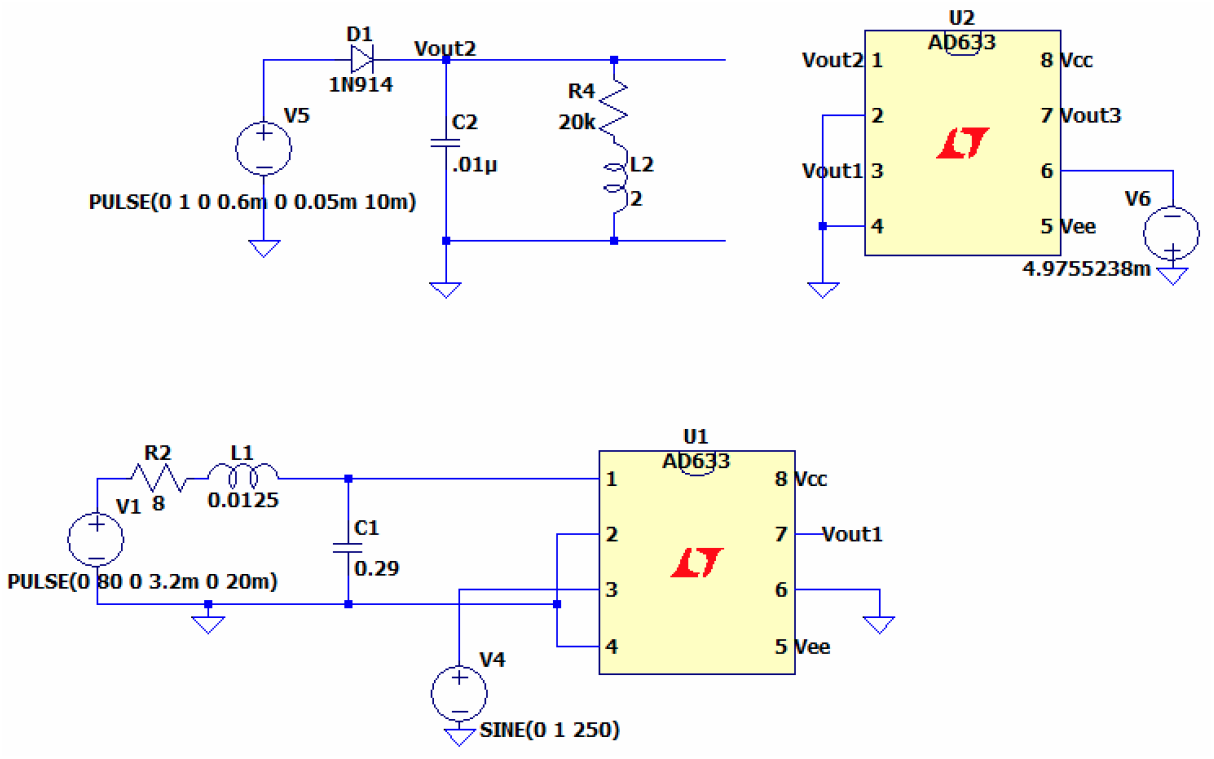
A second AD633 IC is used to produce the convolution of Vout1 and Vout2.

### E. Amplifier and Clamping Circuits

Because the convolution of the responses is very small in magnitude, we designed two operational amplifiers to amplify the latter by 170 times. In the final circuit, we used a third multiplier to pull the initial value of the signal down to the resting potential of –70 mV (Fig. 8).

**Fig. 8.**
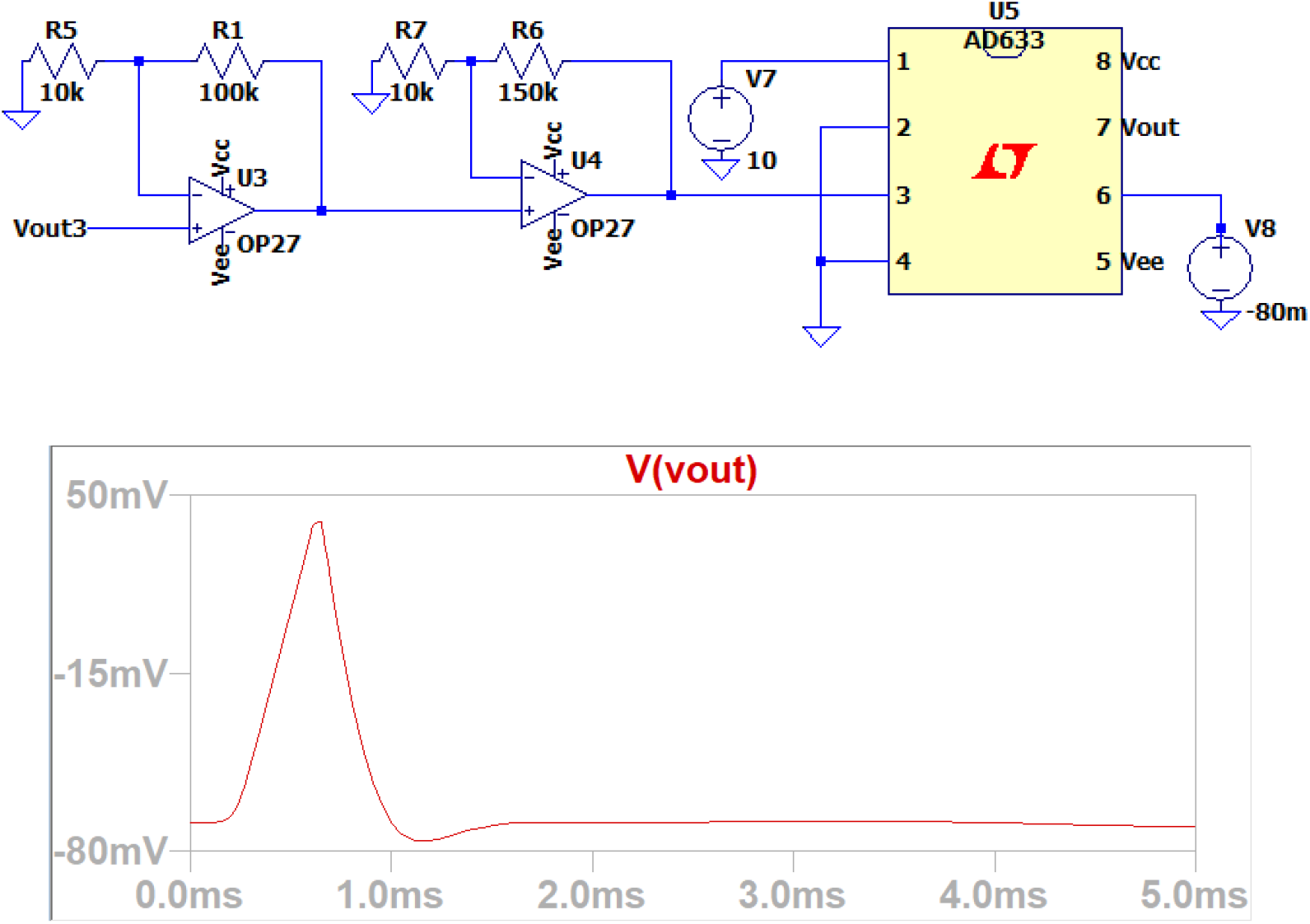
Two OP27 op-amps are set up as noninverting amplifiers. The voltage source at pin 6 of the AD633 multiplier determines the level of shifting down the output signal (pin #7 of the final AD 633 multiplier).

## VII. Summary

The development of an original, quantitative description of the membrane (action) potential displacement *V*_*m*_ was presented in this article. This description is a conductance-based model rooted in cable theory. Unlike the traditional Hodgkin-Huxley model equations of ionic hypothesis, I did not explicitly describe the nerve action potential in the context of ion channels (i.e., the chemistry and physics behind the contribution of different ions to the action potential are not explicit or necessary features of my model).

1. Evidence was presented that three principal factors form a basis on which the membrane potential displacement is described. These three factors are the axon leaky cable conductance, intracellular membrane magnetization, and membrane current modulation.
2. The three hypothesized factors were put into a unified, quantitative form for analytically determining *V*_*m*_.
3. Beginning with substitution of established membrane parameters into the mathematical form, the range of phenomena to which the mathematical form is relevant was demonstrated two ways: (*a*) Computation of the membrane electric field, *E*_*m*_; (*b*) computation of the membrane potential displacement, *V*_*m*_.
4. One of the novelties of this work is that it provides a mechanistic understanding of how intracellular conductance, the thermodynamics of magnetization, and current modulation function together to generate excitation in nerve.
5. Another novel feature of this work is the statistical mechanics description of intracellular magnetization, and how this phenomenon relates to the presence of ions in the membrane channel.
6. The significance of this model is that it offers an original and fundamental advancement in the understanding of the action potential in a unified, analytical description. It provides a conductive, thermodynamic, and electromagnetic explanation of how an action potential propagate in nerve in a single and simple mathematical construct.
7. We synthesized a novel electronic circuit to mimic the electrical bioimpedance of our analytical model. solution. The AD633 multiplier IC is the main component used to produce signals corresponding to the membrane current and voltage and represents the full process of an action potential.

## COMPLIANCE WITH ETHICAL STANDARDS

Conflict of Interests: The authors Robert F. Melendy and Loan Nguyen declare that we have no conflict of interest(s).

## Notes

### Competing Interest Statement

The authors have declared no competing interest.

## REFERENCES

1 R.F. Melendy, Resolving the biophysics of axon transmembrane polarization in a single closed-form description. Journal of Applied Physics, 118(24), (2015).

2 R.F. Melendy, A subsequent closed-form description of propagated signaling phenomena in the membrane of an axon. AIP Advances, 6(5), (2016).

3 A.L. Hodgkin, Evidence for electrical transmission in nerve. Journal of Physiology, 90, 183–210 (1937).

4 J.B. Hursh, Conduction velocity and diameter of nerve fibers. American Journal of Physiology, 127, 131–139 (1939).

5 B. Frankenhaeuser, The ionic currents in the myelinated nerve fiber. Journal of General Physiology, 48, 79–81 (1965).

6 B. Naundorf, F. Wolf, M. Volgushev, Unique features of action potential initiation in cortical neurons. Nature, 440, 1060–1063 (2006).

7 K.S. Cole, H.J. Curtis, Electric impedance of the squid giant axon during activity. Journal of General Physiology, 22, 649–670 (1939).

8 D.E. Goldman, Potential, impedance, and rectification in membranes. Journal of General Physiology, 27, 37–60 (1943).

9 A.L. Hodgkin, B. Katz, The effect of sodium ions on the electrical activity of the giant axon of the squid. Journal of Physiology, 108, 37–77 (1949).

10 J. Koester, S.A. Siegelbaum, in Principles of Neural Science, E.R. Kandel, J.H. Schwartz, T.M. Jessell, Eds. (McGraw-Hill, New York, 2000), pp. 140–149.

11 A.L. Hodgkin, A.F. Huxley, A quantitative description of membrane current and its application to conduction and excitation in nerve. Journal of Physiology, 117, 500–544 (1952).

12 R.E. Taylor, in Physical Techniques in Biological Research, W.L. Natsiik, Ed. (Academic Press, New York, 1963), pp. 219–262.

13 R. Iansek, S.J. Redman, An analysis of the cable properties of spinal motoneurones using a brief intracellular current pulse. Journal of Physiology, 234, 613–636 (1973).

14 W. Rall, J. Segev, The Theoretical Foundation of Dendritic Function: Selected Papers of Wilfrid Rall with Commentaries (MIT Press, Boston, MA, 1995).

15 M. London, C. Meunier, I. Segev, Signal transfer in passive dendrites with nonuniform membrane conductance. Journal of Neuroscience, 19, 8219–8233 (1999).

16 F. Nadim, J. Golowasch, Signal transmission between gap-junctionally coupled passive cables is most effective at an optimal diameter. Journal of Neurophysiology, 95, 3831–3843 (2006).

17 H.M. Lieberstein, On the Hodgkin-Huxley partial differential equation. Mathematical Biosciences, 1, 45–69 (1967).

18 W. Rall, Core Conductor Theory and Cable Properties of Neurons: Handbook of Physiology, the Nervous System, Cellular Biology of Neurons (American Physiological Society, 1977), pp. 39–93.

19 R. West, E. Schutter, G. Wilcox, in The IMA Volumes in Mathematics and its Applications: Evolutionary Algorithms, L.D. Davis et al., Eds. (Springer, New York, 1999), pp. 33–64.

20 C. Bédard, A. Destexhe, A modified cable formalism for modeling neuronal membranes at high frequencies. Biophysical Journal, 94, 1133–1143 (2008).

21 J.J.B. Jack, D. Noble, R.W. Tsien, Electric Current Flow in Excitable Cells (Clarendon Press, Oxford, 1975).

22 D. Sterratt, Principles of Computational Modelling in Neuroscience (Cambridge University Press, Cambridge, 2011).

23 R. Hobbie, Intermediate Physics for Medicine and Biology (AIP Press, New York, 1997).

24 R. Plonsey, R. Barr, Bioelectricity: A Quantitative Approach (Springer, Boston, 2000).

25 N. Sperelakis, N. Sperelakis, Cell Physiology Sourcebook: Essentials of Membrane Biophysics (Academic Press, London, 2012).

26 J. Malmivuo, R. Plonsey, Bioelectromagnetism: Principles and Applications of Bioelectric and Biomagnetic Fields (Oxford University Press, New York, 2000).

27 B. Roth, J. Wikswo, The magnetic field of a single axon: a comparison of theory and experiment. Biophysical Journal, 48, 93–109 (1985).

28 B. Roth, J. Wikswo, The electrical potential and the magnetic field of an axon in a nerve bundle. Mathematical Biosciences, 76, 37–57 (1985).

29 R.S. Wijesinghe, Detection of magnetic fields created by biological tissues. Journal of Electrical and Electronic Systems, 3, 1–7 (2014).

30 B. Greenebaum, F. Barnes, Bioengineering and Biophysical Aspects of Electromagnetic Fields (CRC/Taylor & Francis, Boca Raton, FL., 2007).

31 B. Commoner, J. Townsend, G.E. Pake, Free radicals in biological materials. Nature, 174, 689–691 (1954).

32 V.N. Varfolomeev et al., Paramagnetic properties of hepatic tissues and transplantable hepatomas. Biofizika. 21, 881–886 (1976).

33 R. Pethig, D.B. Kell, The passive electrical properties of biological systems: their significance in physiology, biophysics and biotechnology. Physics in medicine and biology, 32, 933–970 (1987).

34 C. Kittel, Introduction to Solid State Physics (Wiley, New York, 2008).

35 W.T. Coffey, Y.P. Kalmykov, J.T. Waldron, The Langevin Equation, with Applications in Physics, Chemistry, and Electrical Engineering (World Scientific, River Edge, NJ, 1996).

36 J. Koester, S.A. Siegelbaum, in Principles of Neural Science, E.R. Kandel, J.H. Schwartz, T.M. Jessell, Eds. (McGraw-Hill, New York, 2000), pp. 150–169.

37 A.F. Huxley, From overshoot to voltage clamp. Trends in Neurosciences, 25, 553–558 (2002).

38 E.O. Hernández-Ochoa, M.F. Schneider, Voltage clamp methods for the study of membrane currents and SR Ca2+ release in adult skeletal muscle fibers. Progress in Biophysics and Molecular Biology, 108, 98–118 (2012).

39 S.G. Waxman, J.D. Kocsis, P.K. Stys, Eds., The Axon: Structure, Function and Pathophysiology (Oxford University Press, New York, 1995).

40 A.V. Holden, P.G. Haydon, W. Winlow, Multiple equilibria and exotic behavior in excitable membranes. Biological Cybernetics, 46, 167–172 (1983).

41 R. Guttman, S. Lewis, J. Rinzel, Control of repetitive firing in squid axon membrane as a model for a nuroneoscillator. Journal of Physiology, 305, 377–395 (1980).

42 H.R. Leuchtag, Voltage-Sensitive Ion Channels: Biophysics of Molecular Excitability (Springer, New York, Philadelphia, 2008).

43 D.A. Hill, Electromagnetic Fields in Cavities: Deterministic and Statistical Theories (IEEE Press Series on Electromagnetic Wave Theory, NJ, 2009).

44 D.A. McQuarrie, Mathematical Methods for Scientists and Engineers (University Science Books, CA, 2003).

45 R. FitzHugh, Impulses and physiological states in theoretical models of nerve membrane. Biophysical Journal, 1, 445–466 (1961).

46 G. Zhao, Z. Hou, H. Xin, Frequency-selective response of FitzHugh-Nagumo neuron networks via changing random edges. Chaos: An Interdisciplinary Journal of Nonlinear Science, 16, 043107 (2006).

47 S.Y. Gordleeva, et al., Bi-directional astrocytic regulation of neuronal activity within a network. Frontiers in Computational Neuroscience, 6, 104–114 (2012).

48 R.W. Aldrich, P.A. Getting, S.H. Thompson, Inactivation of delayed outward current in molluscan neurone somata. Journal of Physiology, 291, 507–530 (1979).

49 K. Aihara, G. Matsumoto, in Nerve Excitation and Chaos: Dynamical Systems and Nonlinear Oscillations, Gikō Ikegami, Ed. (World Scientific Publishing Co., 1986). Pp. 254–267.

50 J. Rinzel, G. Huguet, Nonlinear Dynamics of Neuronal Excitability, Oscillations, and Coincidence Direction. Communications on Pure and Applied Mathematics, 66(9), 1464–1494 (2013).

51 Morris, H. Lecar, Voltage oscillations in the barnacle giant muscle fiber. Biophysical Journal, 35, 193–213 (1981).

52 T. Sasaki, N. Matsuki, Y. Ikegaya, Action-potential modulation during axonal conduction Science, 331, 599–601 (2011).

53 N.H. Sabah, K.N. Leibovic, The effect of membrane parameters on the properties of the nerve impulse. Biophysical Journal, 12, 1132–1144 (1972).

54 N.F. Britton, Essential Mathematical Biology (Springer-Verlag, London, 2003).

55 J.D. Murray, Mathematical Biology I: An Introduction (Springer-Verlag, Berlin, 2002).

56 E.O. Voit, A First Course in Systems Biology (Garland Science, Taylor & Francis, New York, 2013).

57 R.L. Armstrong, J.D. King, The Electromagnetic Interaction (Prentice Hall, Englewood Cliffs, NJ, 1973).

58 G.B. Arfken, H.J. Weber, F.E. Harris, Mathematical Methods for Physicist: A Comprehensive Guide (Elsevier, MA, 2013).

59 E. Weisstein, CRC Concise Encyclopedia of Mathematics (CRC Press, Boca Raton, 2003).

60 The electrical system of the body: The physics of the nervous system (Medical Physics, University of Notre Dame, n.d., http://www3.nd.edu/~nsl/Lectures/mphysics/).

61 R.I. Macey, in Membrane Physiology, T.E. Andreoli, J.F. Hoffman, D.D. Fanestil, Eds. (Springer, New York, 1980), pp. 125–146.

62 T. Begenisic, Magnitude and location of surface charges on myxicola giant axons. The Journal of General Physiology, 66, 47–65 (1975).

63 J. Enderle, S. Blanchard, J. Bronzino, Introduction to Biomedical Engineering (Elsevier Academic Press, Amsterdam, Boston, London, New York, 2005).

64 P. Smejtek, in Permeability and Stability of Lipid Bilayers, E. Anibal Disalvo, S.A. Simon, Eds. (CRC Press, Boca Raton, Ann Arbor, London, 1994), pp. 197–236.

65 J. Choma, Jr. Electrical Networks: Theory and Analysis. (Wiley Interscience, New York, 1985), pp. 363–411.

66 G.C. Temes, J.W. LaPatra. Circuit Synthesis and Design. (McGraw-Hill, New York, 1977), pp. 51–120.

67 AD633 Integrated Circuit. (2022). Analog Devices. https://www.tme.eu/Document/ce5356ac4efb480c752b9e53289e2634/AD633ARZ-Analog-Devices.pdf

68 LTSpice. (2022). https://Www.Analog.Com/En/Design-Center/Design-Tools-and-Calculators/Ltspice-Simulator.Html. https://www.analog.com/en/design-center/design-tools-and-calculators/ltspice-simulator.html

